# An endodermal subpopulation gives rise to neuromesodermal progenitors in the posterior chick embryo

**DOI:** 10.64898/2026.05.20.726401

**Authors:** Panagiotis Oikonomou, Lisa Calvary, Devany Du, Juni Polansky, Giacomo Gattoni, Connor Lynch, Lingting Shi, Christian Mayer, José McFaline-Figueroa, Nandan L. Nerurkar

## Abstract

Embryogenesis occurs through a progressive narrowing of cell fate potential, initiating with the segregation of three distinct germ layers during gastrulation. Although classically, each germ layer contributes to distinct tissue types as development proceeds, this view has been revised with the discovery of neuromesodermal progenitors (NMPs) — a bipotent progenitor population in the posterior embryo that gives rise to traditionally ectodermal and mesodermal tissues after gastrulation has concluded. However, until now the notion of lineage restriction of the endoderm to gastrointestinal, respiratory, and endocrine tissues has largely remained intact. Here, we describe a unique subpopulation in the chick endoderm that initially lines the ventral surface of Hensen’s node (the amniote organizer). As posterior regression of the node ends with termination of the primitive streak, these cells undergo an FGF-dependent epithelial-to-mesenchymal transition, erasing their endodermal identity as they invade the tailbud and subsequently differentiate into a remarkably broad range of cell types including paraxial, lateral plate, and intermediate mesoderm, and to a lesser extent, notochord and neural tube. Disrupting ingression of node endoderm reduced embryonic axis elongation — a process attributed to mesoderm — by 50%. Through lineage barcoding, single-cell RNA sequencing, and fate mapping experiments, we conclude that the endodermal compartment of Hensen’s node harbors a mixed population of fate restricted and multipotent progenitor cells that give rise to clonal populations spanning traditional germ layer boundaries. These findings illustrate a surprising example of germ layer plasticity and fate convergence across distant progenitor populations during amniote development.

## INTRODUCTION

Gastrulation is a central step in lineage segregation during embryogenesis, with the pluripotent epiblast separating into three distinct germ layers: endoderm, mesoderm, and ectoderm (*1*). While conventionally, it was long accepted that germ layer commitment is exclusively established during gastrulation, this view was revised with the discovery of neuromesodermal progenitors (NMPs), a bipotent cell population that persists beyond gastrulation while maintaining the capacity to generate ectodermal and mesodermal cell types (*2*). In amniote embryos, NMPs derive from the boundary between Hensen’s node and the primitve streak, the caudal lateral epiblast (*3*), and upon ingressing from the epiblast, occupy the chordoneural hinge, the region where the notochord and neural tube transition to the loose, disorganized tissues of the tailbud (*4*, *5*). Molecularly, NMPs are most widely defined by their co-expression of *SOX2* and *TBXT*, classical markers of ectoderm and mesoderm, respectively (*5–7*). However, studies alternatively propose that co-expression of *SOX2* and *TBXT* is not necessary (*8*, *9*) or sufficient (*10*), and that specific activity of the N-1 enhancer regulating expression of *SOX2* is a more appropriate marker of NMPs (*11*, *12*). NMPs have been rigorously studied to understand their embryonic origin (*3*, *13*) and the regulatory logic that drives their commitment to neural and mesodermal fates (*6*, *7*, *14–17*).

While retrospective clonal analyses that first identified NMPs indicated that they do not contribute to gut tissue (*2*), a pair of recent studies suggest that NMPs do contribute to the mouse hindgut (*18*) and its transient, distal most segment, the tailgut (*19*). On the other hand, a role for definitive endoderm in regulating or contributing to NMPs has not been explored. Germ layer plasticity between mesoderm and endoderm has been studied during gastrulation, where the existence of bipotent mesendoderm progenitors has been debated in the amniote epiblast (*20*, *21*). Still, the existence of bipotent mesendoderm progenitors outside the epiblast has not been previously described. Despite insights into the cell behaviors underlying establishment of the definitive endoderm during gastrulation (*22–24*) and later organogenesis of the gut (*25*), much less is known of the intervening stages when the endodermal epithelium occupies the ventral surface of the amniote embryo. As a result, potentially fundamental aspects of endoderm biology, including germ layer plasticity of the definitive endoderm, have been understudied during these stages.

Here, we identify an endodermal subpopulation in the posterior chick embryo that undergoes a previously unreported epithelial-to-mesenchymal transition (EMT), invading the adjacent tailbud mesenchyme from the ventral surface of the embryo and mixing with traditional NMPs, which instead ingress from the epiblast dorsally. Disruption of cell ingression in this endodermal population disrupts axis elongation, a phenomenon primarily attributed to mesodermal tissues (*26*, *27*). Single-cell RNA sequencing (scRNA-seq) and gene expression analyses indicate that prior to ingression, these cells express endodermal markers, but down regulate them upon ingression as they increasingly adopt a broad range of neuromesodermal fates. Through combined lineage barcoding and single-cell transcriptomics, we conclude that the endoderm lining Hensen’s node contains fate restricted progenitors as well as mixed multipotent progenitors that gives rise to clonal populations spanning across traditional germ layer boundaries after gastrulation has ended.

## RESULTS

### The ventral surface of Hensen’s node undergoes an epithelial-to-mesenchymal transition that initiates with the end of node regression

We previously developed an electroporation method that targets high-efficiency transfection to the avian endoderm with no detectable expression in the subadjacent mesoderm (*28–30*). This is apparent from electroporation of the ventral surface of chick embryos with a constitutive GFP reporter at Hamburger Hamilton stage (HH) 10 (36 hours of incubation) (*31*), producing fluorescence that is restricted entirely to the ventral surface by HH15 (Fig. 1A, top). Surprisingly, however, we noted one striking exception: GFP+ cells were dispersed throughout the tailbud mesenchyme of HH15 embryos (Fig. 1A, bottom), appearing to ingress from the ventral surface of the tailbud, but not the neighboring endoderm (Supplemental Fig.S1A) (*28*). The tailbud forms from remnants of Hensen’s node, the amniote organizer, which regresses posteriorly during gastrulation until it reaches the distal end of the primitive streak at HH11 (*32*). By HH18 (embryonic day 3), when posterior endoderm has been internalized to form the hindgut, many GFP+ cells were distributed outside the gut tube along the posterior trunk and tail (Fig. 1B), suggesting that cells lining the ventral surface of the tailbud contribute to non-endodermal tissues. At earlier stages, GFP+ cells were retained within the ventral surface of Hensen’s node, and only began to appear in the neighboring mesenchyme after HH10 (Fig. 1C). Therefore, cells lining the ventral surface of Hensen’s node migrate into the forming tailbud mesenchyme just as the primitive streak terminates and the mode of primary axis elongation shifts from posterior regression of the node toward tailbud outgrowth (*33–35*).

**Figure 1.**
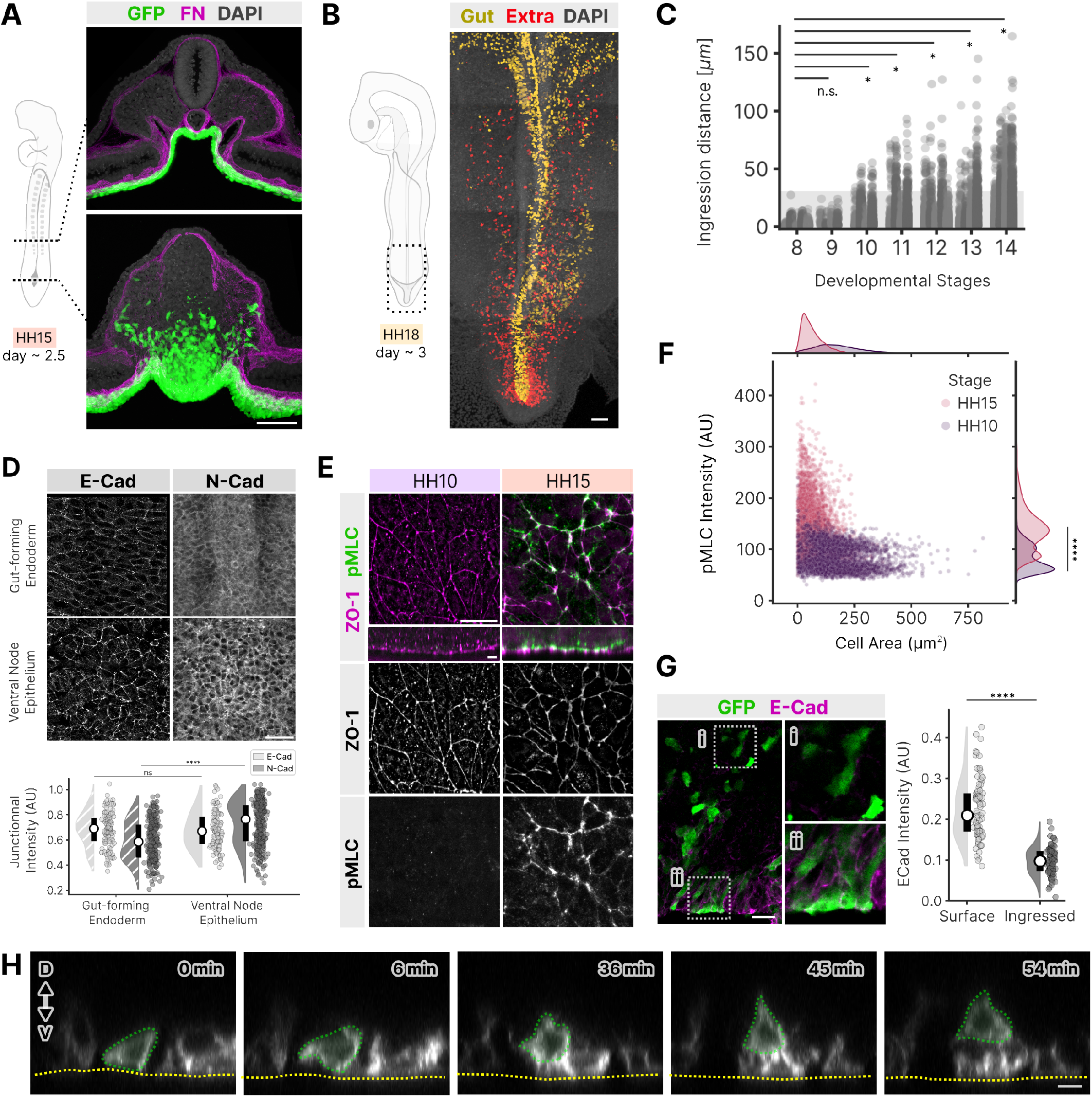
Cells of the ventral node undergo EMT and ingress into the tailbud. (A) Transverse sections of an HH15 chick embryo following ventral epithelium-targeting electroporation with pCAG-GFP, which encodes a constitutive GFP reporter at HH11. Two representative sections are shown along the antero-posterior axis: one anterior to the tailbud (top), the other through the tailbud. Samples counterstained with DAPI (gray) and immunostained for fibronectin (FN). Scale = 50µm. (B) Pseudocolored GFP+ cells (yellow: gut tube, red: extra) in HH18 embryo following electroporation of ventral surface epithelium of pCAG-GFP to label cells at HH11; Scale = 50µm. (C) Distance of cell ingression into the tailbud from the ventral surface of the node/tailbud as a function of developmental stage following pCAG-GFP. Each dot is a single cell, and alternating shades of gray indicate distinct embryos (n=4-5). Transparent gray bar indicates approximate apico-basal height of surface epithelial cells, such that cells appearing beyond this distance have ingressed. One-sided Mann-Whitney U tests, adjusted using the Benjamini-Hochberg False Discovery Rate (FDR). (D) Immunostaining for E-Cadherin and N-Cadherin at the apical surface of the gut forming endoderm and ventral node epithelium at HH15 (top), and quantification of junctional intensity of each (bottom); Scale = 20µm. Student t-test. (E) Immunostaining of phospho-myosin light chain (pMLC) of the ventral node epithelium, counterstained for ZO-1 to visualize apical surface, prior to ingression (HH10) and after onset of ingression at HH15; Scale = 5µm). (F) Quantification of apical cell area and pMLC intensity of single cells in the ventral node epithelium at at HH10 (n = 3633 cells from 8 embryos) and HH15 (n = 5891 cells from 5 embryos). Student t-test. (G) Transverse section of node at HH15 following pCAG-GFP electroporation to label ventral node epithelium at HH10, immunostained for E-Cadherin (left), and quantification of intensity in surface vs. ingressed GFP-expressing cells (right, n = 8919 cells across 5 embryos); Scale = 20µm. (H) Time-lapse stills from *in vivo* imaging of node endoderm during cell ingression (green annotation); apical surface of node epithelium annotated in dashed yellow; Scale = 10µm. n.s. = non-significant, * p *<* 0.05, **** p *<* 0.0001.

To test whether cell ingression from the ventral surface of Hensen’s node constitutes an EMT, we first asked whether the tissue is in fact a polarized epithelium. Indeed, ventral node cells expressed apically localized markers of polarized epithelia, including E-Cadherin (Fig. 1D, Supplemental Fig.S1B) and ZO-1 (Fig. 1E), consistent with the gut forming endodermal epithelium with which the ventral surface is continuous. Junctional intensity of E-Cadherin staining was similar between the ventral node epithelium (VNE) and the neighboring gut-forming endoderm. However, once ingression was underway at HH15, N-Cadherin was significantly increased at cell junctions in the VNE population (Fig. 1D), consistent with the onset of EMT. This was further supported by significant upregulation of apically localized phospho-myosin light chain (pMLC) at HH15 compared to HH10 (Fig. 1E, F, Supplemental Fig.S1C), a decrease in mean apical surface area, and the emergence of an inverse correlation between junctional pMLC intensity and apical surface area at HH15 compared to HH10 (Fig. 1F, Supplemental Fig.S1D). These data suggest cell ingression is correlated with apical constriction, another hallmark of EMT (*36*, *37*). Ingressed GFP+ cells down-regulated E-Cadherin relative to cells at the ventral surface (Fig. 1G), indicating a loss of epithelial identity upon ingression. Finally, VNE ingression was confirmed directly by live imaging (Fig. 1H, Supplemental Movie 1), with cells losing their contacts with the apical surface as they migrate basally into the neighboring mesenchyme. Together, these data suggest that the ventral surface of Hensen’s node comprises an epithelial cell population that initiates EMT once node regression completes and the tailbud begins to form.

### The ventral node epithelium (VNE) is endoderm

Localized EMT within the VNE raises the question of what the identity and origin of these cells are. Previous fate mapping studies in the chick demonstrated that between HH3 and HH4, the ventral surface of Hensen’s node is populated by definitive endoderm, cells deriving from the epiblast that ingress through the anterior primitive streak and re-epithelialize at the ventral surface of the embryo (*38–40*). We found that VNE cells tagged at HH3 were displaced posteriorly with the node as it regresses through HH10 (Fig. 2A). Early during node regression, VNE cells expressed the classical definitive endoderm marker SOX17 (Fig. 2B). However, by HH10, SOX17 was diminished throughout the posterior endoderm, including both the presumptive hindgut and the node region (Fig. 2B). Taken together, we conclude that the VNE is endodermally derived, but downregulates SOX17 prior to the onset of EMT.

**Figure 2.**
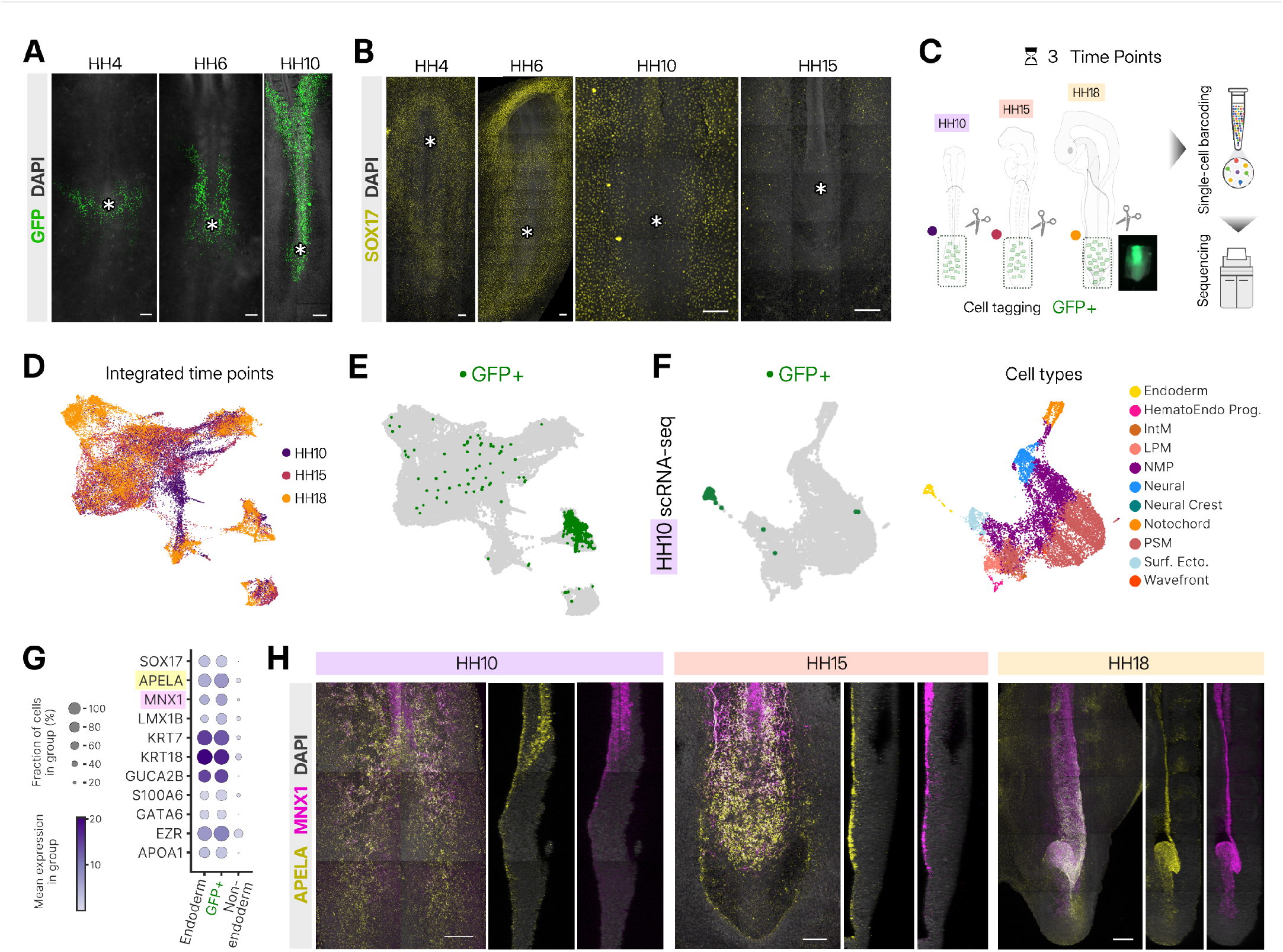
The ventral node epithelium is an endodermal tissue. (A) Fate mapping by *3nls-GFP* RNA electroporation to tag node epithelium during gastrulation at HH3 and imaged through the end of gastrulation at HH10; * denotes location of Hensen’s node; Scale = 100µm. (B) Ventral view of HH4-15 embryos immunostained in whole mount to visualize SOX17; * denotes location of Hensen’s node; Scale = 100µm;. (C) Overview of study design: embryos were electroporated with pCAG-GFP to tag the posterior endoderm, then tissue was collected at HH10, HH15, and HH18 for single-cell RNA sequencing. (D) PaCMAP embedding of the batch-integrated dataset across developmental stages. (E) Cells with detectable GFP transcripts indicated in green on batch-integrated PaCMAP embedding of batch the integrated datasets. (F) PaCMAP embedding of HH10 scRNA-seq dataset, with GFP-expressing cells indicated in green (left) and cell type annotations as indicated (right). (G) Dot plot representation of a subset of the top 20 differentially expressed genes between the endoderm cluster and all other cell types. (E) Hybridization chain reaction (HCR) for multiplexed detection of *APELA* (yellow) and *MNX1* (magenta) expression at HH10 (left), HH15 (center) and HH18 (right); Scale = 100µm.

To establish the identity of VNE and its ingressed descendants in the full cellular context of posterior chick development, we performed single-cell RNA sequencing (scRNA-seq). Embryos were electroporated with a constitutive GFP reporter to tag the ventral epithelium and its descendants, and all posterior tissues were collected for scRNA-seq at HH10, HH15, and HH18 (Fig. 2C). Batch integration yielded an embedding with smooth transitions across developmental stages (Fig. 2D, Supplemental Fig.S2A). Detection of GFP transcripts was used to identify VNE descendants within the broader dataset (Fig. 2E, Supplemental Fig.S2B). Focusing first on HH10, prior to the onset of ingression, the endoderm was among the most uniform and transcriptionally distinct clusters in PaCMAP space (Fig. 2F, Supplemental Fig.S3A). 97% of GFP-expressing cells were embedded within the endoderm cluster at HH10 (Fig. 2F, Supplemental Fig.S3A), and GFP-expressing cells expressed top endoderm-specific differentially-expressed genes (DEGs, Fig. 2G, Supplemental Fig.S4). Therefore, despite downregulation of SOX17, a transcriptome-wide view of cell identity confirms that the VNE is endoderm at the onset of cell ingression. Guided by endoderm-specific DEGs, we determined by in situ hybridization chain reaction (HCR) that co-expression of *APELA* (*19*) and *MNX1* (*41*, *42*) serves as a faithful and persistent marker of the posterior chick endoderm through HH18 (Fig. 2H), when SOX17 is no longer detected. Consistent with their endodermal identity, VNE strongly coexpressed *APELA* and *MNX1* (Fig. 2H). Moving forward, we refer to this cell population as node endoderm (NE).

### Node endoderm (NE) gives rise to neuromesodermal progenitor cells (NMPs)

Expression of endodermal markers *APELA* and *MNX1* is restricted to the ventral surface of the embryo at HH15 (Fig. 2H), while GFP+ NE descendants extend deeper into the tissue (e.g. Fig. 1A). This suggests that NE cells may lose their endodermal identity upon ingression into the tailbud, raising the question of what fate these cells ultimately adopt. At the transcriptome level, GFP+ cells were more broadly distributed in the PaCMAP embedding at HH15 (Fig. 1A-C), including many cells clustered with NMPs and paraxial mesodermal fates (Fig. 3A). Immunostaining for SOX2 and TBXT revealed that 71.7%*±*11.5 of GFP+ cells (n=506 cells from 8 embryos) in the tailbud were double positive for these markers (Fig. 3B), compared to 0.0% of GFP+ cells anterior to the tail bud (n=150 cells from 5 embryos). Activity of the *SOX2* enhancer element N-1, which serves as an alternative marker for NMPs (*12*, *43*), was detected within the NE and posterior-most gut forming endoderm, but significantly reduced throughout the surrounding endoderm (Supplemental Fig.S5). Together, these data suggest that NE and its ingressed descendants adopt an NMP identity.

**Figure 3.**
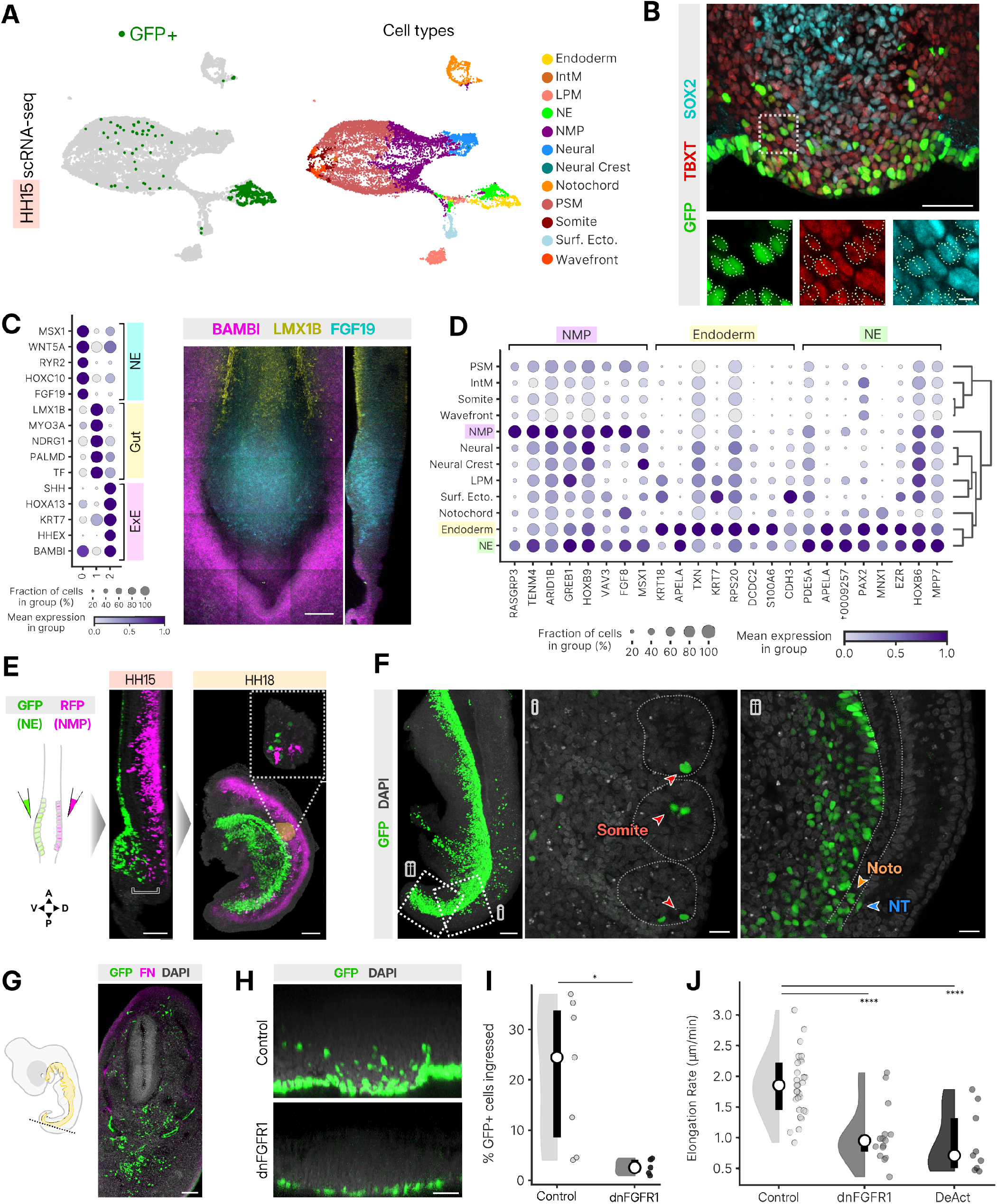
Node Endoderm gives rise to NMPs. (A) PaCMAP embedding of HH15 scRNA-seq dataset, with GFP-expressing cells indicated in green (left) and cell type annotations as indicated (right). (B) Transverse section of node at HH15 following pCAG-3nls-GFP electroporation to label ventral node epithelium at HH10, immunostained for TBXT (red) and SOX2 (cyan); boxed region magnified below with individual nuclei outlined to appreciate colocalization; Scale = 50µm (top) and 5µm (bottom). (C) Differentially expressed genes (left) within three subpopulations of the posterior HH15 endoderm: node endoderm (NE), gut forming endoderm (Gut), and extraembryonic endoderm (ExE). Multiplexed HCR to visualize expression of *FGF19* (cyan), *LMX1B* (yellow), and *BAMBI* (magenta) to demarcate NE, Gut, and ExE subpopulations, respectively; Scale = 100µm. (D) Hierarchical clustering and differential gene expression of cell types in the HH15 scRNA-seq dataset. (E) Dual electroporation targeting pCAG-GFP to the endoderm ventrally and pCAG-myrRFP to the caudal lateral epiblast dorsally at HH9-10 (schematic, left), visualized in optical sagittal sections of optically cleared embryos developed to HH15 (center) and HH18 (right) and counter-stained with DAPI (gray). Inset: magnified view of somite containing both GFP+ and RFP+ cells; A/P and D/V denote anterior/posterior and dorsal/ventral embryonic axes; square bracket indicates the chordoneural hinge (Supplemental Fig. S1A); Scale = 100µm. (F) 3D volumetric imaging of optically cleared HH18 embryo following electroporation of posterior endoderm with pCAG-H2B:GFP at HH11, counter-stained with DAPI (gray). Magnified optical sections isolated from boxed regions i (center) and ii (right) to visualize isolated GFP+ cells in somites, tail notochord (Noto), and neural tube (NT); Scale = 100µm (left), 20µm (center, right). (G) Transverse view of embryonic day 5 embryo following *in ovo* electroporation of posterior endoderm with pCAG-GFP, counter-stained for fibronectin (FN, magenta) and DAPI (gray). Scale = 100µm. (H, I) Electroporation of pCAG-GFP (Control) vs. dnFGFR1-IRES-GFP at HH11 inhibits NE ingression (H), quantified in (I). Scale = 20µm. (J) Axis elongation rate quantified between HH10 and HH14 following electroporation of the NE with pCAG-GFP (Control, n=29), dnFGFR1-IRES-GFP (n=15), or pCAG-DeAct-GFP (n=9). * p *<* 0.05, **** p *<* 0.0001.

Widespread EMT and expression of NMP markers in the NE suggests that this population is fundamentally distinct from traditional endoderm, which folds into the gut tube and generates the inner epithelial lining of respiratory, endocrine, and gastrointestinal organs. To investigate this apparent heterogeneity, we examined transcriptional variation within the posterior endoderm of HH15 embryos, identifying three distinct subpopulations (Fig. 3C, Supplemental Fig.S6A, B): hindgut-forming endoderm that is *LMX1B*+, anterior to the node, extraembryonic endoderm that is *BAMBI*+, posterior and lateral to the node, and NE that is *FGF19*+ (Fig. 3C). Several additional genes were confirmed by HCR to be expressed in the NE, but undetected in the neighboring gut and extramebryonic endoderm, including *FGF8*, *TBXT*, *SLIT2*, *CDX2*, and *MSX1* (Supplemental Fig.S6C). Recognizing NE as separable from gut forming and extraembryonic endoderm, we refined our cell type annotations (Table S1) to designate NE cells as distinct cell type. Notably, *SOX2* and *TBXT* were co-expressed in both NMPs and this newly designated NE cluster at HH15 (Supplemental Fig.S6D), despite neither gene being used among the markers for annotating the NE cell type. Hierarchical clustering and identification of top DEGs revealed that NE most closely aligned with gut-forming endoderm, but also highly expressed NMP-specific genes (Fig. 3D).

When dual-labeling experiments were performed, targeting expression of GFP to NE cells ventrally and RFP to the caudal lateral epiblast where traditional NMPs emerge dorsally, we found that the two populations converge on the chordoneural hinge by HH15 (Fig. 3E), and at later stages, descendants of both NE and traditional NMPs incorporated into mature somites together (right inset, Fig. 3E). When NE was labeled by electroporation at the onset of ingression at HH11 and embryos were cultured to HH18, GFP+ descendants were observed in somites, paraxial mesoderm, and, in rare cases, notochord and neural tube as well (Fig. 3F, Supplemental Fig.S7A). Sparse *in ovo* electroporation of NE for long-term labeling revealed that GFP+ NE descendants remain abundant throughout mesodermal derivatives including the sclerotome and dermomyotome of embryos at embryonic day (E5), when apoptosis of tail cells is complete (*44*) and organogenesis is well underway (Fig. 3G). This suggests a sustained role for NE descendants in posterior development as more mature cell types and organs begin to form.

In sum, the endodermal compartment of Hensen’s node harbors a previously unappreciated source of NMPs with a strong bias toward mesodermal fates. This is supported by single-cell transcriptional states (Fig. 3A, D), co-expression of SOX2/TBXT by NE descendants (Fig. 3B, Supplemental Fig. S6D), activity of the N-1 enhancer of *SOX2* (Supplemental Fig.S5), convergence of NE descendants with traditional NMPs on the chordoneural hinge (Fig. 3E), and later contribution of NE descendants to mesodermal, and to some extent neural, fates (Fig. 3E-G,Supplemental Fig.S7A).

### FGF-dependent cell ingression from the node endoderm is required for axis elongation

To test the functional importance of NE as an NMP source, we aimed to block cell ingression and observe the effects on posterior development. Given the central role of FGF signaling in regulating many aspects of posterior development (*28*, *45–47*), and its requirement for mesoderm induction and EMT during gastrulation (*48–50*), we tested whether FGF signaling is required for ingression of NE as well. FGF activity was detected throughout the NE (Supplemental Fig.S7B), and suppression of FGF signaling via misexpression of a dominant negative mutant of the receptor FGFR1 (dnFGFR1, Supplemental Fig.S7B, C) (*51*) inhibited NE cell ingression (Fig. 3H, I). Remarkably, suppression of FGF-dependent ingression in the NE resulted in a marked reduction in axis elongation rate (Fig. 3J). Ablation of NE via targeted electroporation with DeAct-SpvB, a construct that severely disrupts the actin cytoskeleton (*30*, *52*), produced a similar reduction in axis elongation rate to dnFGFR1 misexpression, suggesting that FGF signaling is a major driver of NE ingression. Together, these results suggest that NE plays a significant role in axis elongation, although it is not yet clear whether this is directly as a source of mesodermal cells that expand the antero-posterior axis, or indirectly via cell-cell communication or other interactions downstream of FGF signaling within the NE. Nonetheless, these findings suggest a previously unappreciated role for definitive endoderm in elongation of the primary body axis, a fundamental morphogenetic event that has primarily been attributed to mesoderm (*1*, *26*, *27*).

### The node endoderm is a mixed multipotent progenitor population that gives rise to clonal populations across germ layers

The above findings demonstrate that cells originating with the NE adopt a range of fates. However, how fate decisions are made by individual cells within the NE is unclear. As a first step in this direction, we interrogated inferred lineage trajectories and cell potency in the integrated single-cell dataset comprising posterior tissues of the HH10 (at the onset of ingression), HH15 (ingression well underway), and HH18 embryo (the latest developmental stage accessible via *ex ovo* EC culture, Fig. 2C). GFP-expressing descendants of NE achieved progressively diversified fates from HH10 to HH18, although many cells retained their endodermal identity at HH18, including both gut-forming and node endoderm (Fig 4A, Supplemental Fig. S3). Multipotency was estimated via CytoTRACE2, a deep learning tool trained on diverse developmental scRNA-seq datasets to identify developmental potential and differentiation landscape (*53*). CytoTRACE2 potency scores (Supplemental Fig. S8A) revealed a complex mixture of differentiation potential across the dataset, and identified the transcriptional state of NE as most consistent with a multipotent progenitor population (Fig 4B). This is notable given that marker genes used to annotate the NE (Table S1) do not themselves reflect common pluripotency genes that dictate CytoTRACE2 cell classification.

**Figure 4.**
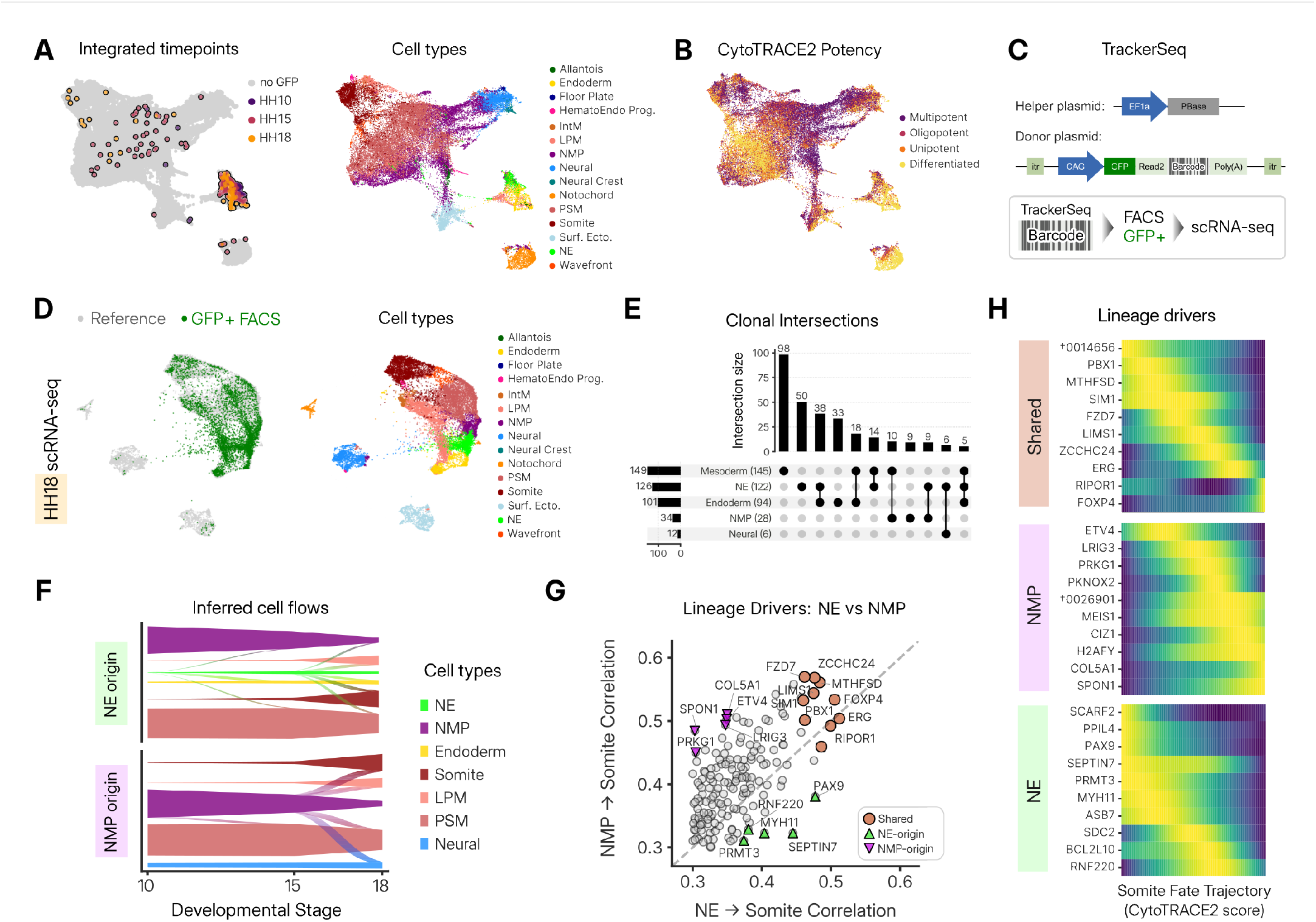
Lineage tracing and trajectory inference. (A) GFP-expressing cells indicated by developmental stage on PaCMAP embedding of scRNA-seq dataset integrated across HH10, HH15, and HH18 datasets (left), and associated cell type annotations (right). (B) Inferred cell potency states associated with cell types as indicated in (A, right), computed from CytoTRACE2 potency scores (Supplemental Fig. S8A). (C) TrackerSeq plasmid design and experimental workflow: posterior endoderm co-electroporated at HH10 with a Donor plasmid library containing unique barcode sequences downstream of GFP and a constitutive CAG driver, flanked by itr sequences, together with a Helper plasmid for genomic integration of the Donor construct. At HH18, cells from posterior chick embryo were dissociated and GFP-expressing cells enriched by FACS prior to scRNA-seq to recover lineage barcodes together with transcriptomes. (D) FACS-enriched GFP-expressing cells (endoderm descendants) registered against the unsorted HH18 scRNA-seq dataset (left), in order to define cell types as annotated in the latter (right). (E) TrackerSeq-based UpSet plot indicating the top 11 clonal distributions of lineage relationships. Vertical columns indicate intersection size (number of clonal populations for a given distribution of cell fates), while rows indicate number of clones associated with a particular lineage. Filled circles connected by a line indicate shared clonality across the spanned lineages. For a more resolved distribution of lineages across all clonal populations detected, see Supplemental Fig. S8D. (F) Lineage trajectories inferred from scRNA-Seq datasets across HH10, HH15, and HH18, constructed via optimal transport to identify lineage trajectories connecting two progenitor populations at HH10, NE (top) and NMPs (bottom), to downstream cell types at HH18. (G) Correlation plot for putative lineage drivers associated with inferred trajectories from NMPs (y-axis) and NE (x-axis) toward somite fates, where shared drivers are associated with both progenitor types, and NMP-origin and NE-origin indicated putative drivers that are selectively enriched among NMP and NE progenitors, respectively. (H) Expression dynamics of shared, NMP-associated, and NE-associated lineage drivers (top 10 highest correlations in G) ordered by peak expression along the inferred somite fate trajectory (CytoTRACE2 potency score).

While inference tools such as CytoTRACE2 can be highly informative for interrogating putative differentiation states from existing datasets, true lineage requires tracking heritable signatures across progeny of single cells. To do this, we utilized TrackerSeq, a lineage barcoding technique compatible with scRNA-seq, so that lineage and cell states can be simultaneously determined (*54*). The posterior endoderm was co-electroporated at HH10 with a plasmid library encoding GFP upstream of one of 125 million possible unique barcode sequences, together with a helper plasmid for transposase-mediated barcode integration (Fig 4C). At HH18, tailbuds were dissociated and FACS-enriched for GFP+ cells, followed by scRNA-seq. Cell types were determined by registering sorted cells against the HH18 reference dataset, which reflects the full distribution of endodermal and non-endodermal cell types in the posterior chick embryo (Fig. 4D). Because this dataset was FACS-enriched for NE descendants, we observed a broader range of cell identities, including cells distributed across all three germ layers. Consistent with the reference single-cell dataset (Fig 4A), endoderm-derived cells were enriched within the NE, gut-forming endoderm, and NMP clusters, but were also abundantly distributed across a diversity of mesodermal lineages, including paraxial, lateral plate, and intermediate mesoderm (Fig. 4D, Supplemental Fig.S8B). Though less frequent, endoderm descendants were also associated with notochord, surface ectoderm, and neural tube identities as well (Fig. 4D). Analysis of barcode distributions (Supplemental Fig.S8C) revealed that the most abundant clonal populations were restricted to mesoderm (Fig. 4E), even though barcodes were delivered at HH10, when NE cells are decidedly of endodermal identity (Fig. 2F, Supplemental Fig.S3A, S4). This suggests that NE likely contains a large proportion of unipotent progenitors committed to mesodermal fates even prior to the onset of EMT. On the other hand, several clonal populations were identified that spanned multiple cell types, even crossing conventional germ layer boundaries (Fig. 4E, Supplemental Fig.S8D). Clonal populations were detected that spanned NE and either endoderm, mesoderm, NMPs, or ectoderm (though this fraction was low). These findings further support the existence of fate-restricted progenitors in the NE that are capable of self-renewal and differentiation into one of the three germ layers. However, we also observed clonal populations that spanned endoderm and mesoderm — but not NE — supporting the existence of bipotent mesendoderm-like cells within the NE as well. Taken together, these data indicate that NE contains a mixed multipotent population of fate-restricted and bipotent progenitor cells.

### Fate convergence of NE and NMP via distinct inferred trajectories

The shared contribution of NE and traditional NMPs to mature somites (Fig. 3, 4, and Supplemental Fig.S7A, S8D) constitutes a surprising example of fate convergence across germ layers. To compare how such dissimilar progenitors converge on the same fate, we used optimal transport (*55*, *56*) to construct inferred differentiation trajectories, mapping between scRNA-seq datasets at HH10, HH15, and HH18. Both NE and conventional NMPs were predicted to give rise to somites based on transport of cell states across developmental stages (Fig. 4F). Computing putative lineage drivers for both progenitor populations returned many genes previously implicated in somitogenesis, including *FOXC2* (*57*), *FZD7* (*58*), and *PBX1* (*59*) (Fig. 4G). However, we also identified genes that — to our knowledge — have not previously been implicated in somitogenesis, including *FOXP4* and *ZCCHC24* (Fig. 4G). Of the many putative lineage drivers identified, several were differentially enriched along either NE or NMP trajectories of somite differentiation (Fig. 4G), suggesting that fate convergence may occur via progenitor cell-type specific mechanims. Lineage drivers shared by both progenitor populations displayed a sequential cascade of gene expression along the trajectory of somite fate specification (Fig. 4H). On the other hand, progenitor-specific lineage drivers had distinct signatures, with most NE-specific genes activated early and NMP-specific drivers enriched late (Fig. 4H). Early activation of NE-specific genes may reflect a transcriptional program associated with germ layer plasticity, as cells suppress their endodermal identity and transition toward paraxial mesoderm. It is not yet clear whether late activation of NMP-specific drivers reflects refinement of the somite program that is not required of NE-descendants or, alternatively, that these reflect broader functional differences between NE and NMP derived paraxial mesoderm cells. Together, these data illustrate the transcriptional dynamics associated with fate convergence across distant progenitor populations and reveal potential insights into germ layer plasticity in the posterior chick embryo.

## DISCUSSION

The present study reveals a previously unappreciated source of progenitor cells in the posterior endoderm that give rise to diverse cell types spanning across traditional germ layer boundaries. Cells that initially form the endodermal compartment of Hensen’s node undergo EMT from the ventral surface of the embryo at the onset of tailbud formation (Fig. 1), transitioning from an endodermal identity (Fig. 2) toward a neuromesodermal progenitor state upon ingression (Fig. 3). Lineage barcoding suggests the node endoderm harbors a mixture of bipotent and fate-restricted unipotent progenitors that generate cells across endoderm, mesoderm, and to a lesser extent, ectodermal fates (Fig. 3, 4). While the present work is limited to the chick embryo, a subpopulation of *SOX2/TBXT* double positive cells that also express endodermal markers is apparent in published mouse (*60*) and macaque (*61*) datasets at equivalent developmental stages (Supplemental Fig.S9), suggesting that the role of node endoderm as a source of neuromesodermal progenitors may be conserved among amniotes more broadly.

The existence of mesendoderm cells in the amniote embryo has been disputed in the literature, with some arguing that a persistent population in the epiblast is capable of generating endodermal and mesodermal cells (*20*), while others suggest that fate restricted endoderm and mesoderm progenitors are spatially mixed within epiblast, but a true mesendodermal state is brief and transient (*21*, *24*, *62*). Here, we find through lineage barcoding that the node endoderm does contain true bipotent ‘mesendoderm’ cells capable of generating endodermal and mesodermal fates from a single parent cell. However, these cells are distinct from the conventional notion of mesendoderm, because they arise from definitive endoderm after gastrulation, rather than from the pluripotent epiblast during gastrulation.

Despite the broad range of cell types generated from node endoderm, our lineage barcoding experiments identified only a single clonal population that spanned all three germ layers, suggesting that few, if any, NE cells retain the ability to generate endoderm, mesoderm, and ectoderm. Limitations to transfection efficiency *in ovo*, along with limitations in the developmental stages achievable via *ex ovo* culture (*63*), limited our lineage barcoding experiments to the developmental window spanning HH11 to HH18. Notably, however, cells retaining a node endoderm identity persisted through HH18, and were predicted to be multipotent by CytoTRACE2 (Fig. 3B). This suggests that at HH18, progenitors within node endoderm are not yet exhausted, and it is possible that longer term *in ovo* lineage tracing experiments may reveal a broader range of clonally linked cell fates.

The present study further complicates an ongoing question of what defines an NMP (*64*, *65*), demonstrating an endodermal source of NMPs that is similar to traditional NMPs, with some important differences. For example, NE cells are marked by *SOX2*/*TBXT* co-expression and SOX2 enhancer N-1 activity, and give rise to cells that adopt paraxial mesoderm and neural tube fates, but do not derive from the caudal lateral epiblast as tradtional NMPs do. We find here that FGF signaling is necessary for NE cells to undergo EMT and contribute to axis elongation (Fig. 3G-I), whereas studies in zebrafish indicate that for traditional NMPs, FGF signaling is required to terminate EMT instead (*66*). We observed in the distal tailbud of HH18 and later embryos that descendants of NE are incorporated into the notochord, while similar contributions have not been observed for NMPs (*2*, *7*, *67*). We find via lineage barcoding that the NE contains bipotent progenitors capable of making gut endoderm and mesoderm. While there is some evidence that cells of the SOX2/TBXT double-positive lineage contribute to the distal gut in mice (*18*), there is no evidence of such bipotent progenitors within the traditional NMP population as yet. *In vitro* models for NMP biology are strongly biased toward neural rather than mesodermal fates (*8*, *68*), and *in vivo* studies suggest a significant portion of SOX2/TBXT expressing cells are committed exclusively to neural fates (*10*). Conversely, we find that NE descendants are strongly biased toward mesodermal fates (Fig. 4D). Indeed, the significant role of NE in axis elongation *in vivo* suggests that *in vitro* and organoid NMP models that do not include an endodermal cell population may miss fundamental aspects of *in vivo* development of the posterior embryo. From the transcriptional standpoint, NE co-expresses many of the genes enriched in NMPs (Fig. 3D), yet trajectory inference revealed differences in the predicted lineage drivers that underlie fate convergence from these two progenitor populations towards paraxial mesoderm differentiation (Fig. 4G,H). Prior studies have noted heterogeneity among NMPs — defined by co-expression of *SOX2* and *TBXT* — *in vivo* (*8*, *18*, *69*). In light of the present work, it is possible that this heterogeneity reflects the disparate embryonic origins of SOX2+/TBXT+ cells that have not previously been appreciated.

In summary, we identify the node endoderm as a novel progenitor population in the posterior chick embryo that erases its germ layer identity before contributing to a remarkably broad range of cell types across the endoderm, mesoderm, and ectoderm, representing a surprising example of cell plasticity and fate convergence across distant progenitor populations. These findings contribute to the emerging view that many ‘classical’ principles of development, established through extensive studies of gastrulation and early fate determination, may not fully capture the complexity of cell fate determination during posterior development of the vertebrate embryo.

## METHODS

### Chicken embryology techniques, electroporations

Fertilized White Leghorn chicken (Gallus gallus domesticus) eggs were acquired from the University of Connecticut Poultry Farm and incubated at 37° C and 60% humidity to the desired stage (*31*). For *ex ovo* manipulation, embryos were harvested onto filter paper rings and transferred to plates containing EC culture medium (*63*). Endoderm electroporation was carried out as previously described (*28*, *29*), with the embryo placed atop a square 2mm x 2mm electrode, submerged in PBS, and 3.5 µg/µL plasmid DNA solution in 5% sucrose and 0.1% Fast Green FCF applied directly to the ventral surface. A second square electrode was placed over the embryo and electroporation pulses applied using a Nepa21 electroporator (Nepa Gene, Ichikawa City, Japan). For dual electroporations targeting the node endoderm and caudal lateral epiblast, embryos were harvested at HH10, and after electroporating the ventral surface to target pCAG-GFP to the node endoderm as described, the embryo was flipped to expose the dorsal surface, and a pulled glass capillary needle was inserted between the epiblast and vitelline membrane to deliver pCAG-myrRFP to the caudal lateral epiblast. FGF activity was detected using a live FGF reporter comprising the FGF-responsive DUSP6 promoter upstream of a nuclear-localized mScarlet, co-electroporated with a constitutive nuclear-localized BFP reporter (*28*). Similarly, activity of the N-1 enhancer associated with SOX2, was detected by co-electroporation of the reporter along with pCAG-H2B-GFP. Ablation experiments were performed by electroporation a plasmid encoding DeAct-SpvB, an ADP ribosyltransferase that massively disrupts F-actin, fused to GFP (*30*, *52*); to limit the range of transfection, these electroporations were performed with plasmid DNA in 10% sucrose to limit diffusion of DNA from the site of delivery. Because DNA electroporation efficiency was limited in early endoderm, HH3 fate maps were instead generated using RNA electroporation: *3nls-GFP* mRNA was generated by *in vitro* transcription of linearized pCS2-3nls-EGFP (Addgene #165400) and electroporated at 500 µg/µL mRNA in 10 % sucrose, and 0.1 % fast green as previously described (*30*). For longer term fate mapping experiments that extended beyond HH18, the upper limit of EC culture, *in ovo* endoderm electroporations were performed. Briefly, eggs were windowed at HH10/11, and plasmid DNA injected between the yolk and the ventral epithelium, followed by placement of wire electrodes atop the vitelline membrane and in the yolk below. Axis elongation rate was quantified from bright field images by manually measuring the change in distance between somite 8 and the caudal extent of the embryo from the time of electroporation to 16 hours later.

### Immunofluorescence, in situ Hybridzation Chain Reaction (HCR), and live imaging

For immunostaining, embryos were fixed overnight at 4°C in 4% paraformaldehyde in PBS, rinsed, and dissected free of surrounding extraembryonic tissue and filter paper. For whole mount staining, embryos were washed (PBS with 0.5% Triton was used for all washes) and incubated in blocking solution (10% heat inactivated goat serum) at room temperature for 2 hours, then incubated overnight at 4°C with primary antibody in pre-block buffer (1%DMSO, 3% BSA diluted in PBS with 0.1% Triton) at the following dilutions: Fibronectin (DSHB, B3/D6; 1:200), SOX17 (R&D Systems AF1924; 1:400), ZO-1 (Invitrogen 33-9100; 1:250), pMLC (Cell Signalling 3671; 1:100), E-Cadherin (BD Bioscience 610181; 1:100), N-Cadherin (BD Bioscience 610920; 1:100), Anti-GFP (Abcam ab13970; 1:1000). Antigen retrieval was performed by 1 hour incubation at 65°C in citrate buffer to improve antibody binding for E-Cadherin, N-Cadherin and pMLC. After 5 washes of 5 minutes each, embryos were incubated with secondary antibodies and DAPI (Invitrogen D1306; 2 µg/mL) overnight at 4°C, washed, and mounted on glass slides for imaging in the optical clearing agent RapiClear (RapiClear 1.49, Sunjin Labs). A subset of fixed embryos were vibrotomed prior to staining, by embedding in 4% low-melt agarose and collecting serial 100 µm slices in either transverse or sagittal orientations using a Leica VT 1200 S vibratome. For section immunofluorescence, fixed samples were incubated in increasing concentrations of sucrose before embedding and freezing in O.C.T., following by sectioning to 16 µm (Leica 3050S cryostat), and following antigen retrieval in pre-warmed citrate buffer (pH=6) for 10 minutes in a vegetable steamer, immunostained as described previously (*28*), with the following dilutions: GFP (above), E-Cadherin (above), TBXT (R&D Systems, AF2085-SP; 1:300), and SOX2 (MilliporeSigma, AB5603; 1:150). For SOX2 and TBXT, 1% DMSO was included in the blocking buffer for blocking and antibody incubation.

HCRs were performed per manufacturer’s instructions (Molecular Instruments). Briefly, embryo tailbuds were dissected from the surrounding extraembryonic tissue, fixed for 4 hours at room temperature in 4% paraformaldehyde in PBS, and rinsed in PBS with 0.5% Triton. After serial de-(and re-)hydration in MetOH, samples were prehybridized in probe hybridization buffer and then incubated overnight with probes (10 nM) and buffer at at 37°C. The next day, samples were pre-amplified and then incubated overnight at room temperature in hairpin (115 nM) with DAPI (2 µg/mL). After SSCT and PBS Triton washes, embryos were cleared in RapiClear (RapiClear 1.49, Sunjin Labs) and mounted on glass slides for imaging. Fluorescence imaging was performed on an inverted Zeiss LSM 880 confocal microscope, coupled with a Coherent Ultra II two-photon laser. Live imaging was performed in Mat-Tek glass bottomed dishes with an environmental chamber maintain 37.5°C (*28*).

### Image analysis and Data processing

Image analyses on 3D cleared volumes were performed using the open-source image visualization Python library Napari (*70*), preprocessing functions from the sci-kit image library (*71*), and the cell segmentation algorithm Cellpose (*72*). After cell segmentation and (ventral) surface extraction of the endoderm (*29*), a euclidean distance transform was calculated to compute ingression depth of each cell centroid. E-Cadherin and N-Cadherin junctional intensities were quantified using the segmentation mask (3 pixel width) in regions of 20 µm^2^in the center of the node of 200 µm (gut-forming endoderm). To avoid variability between samples, each sample’s intensity was normalized to its top 99.5th percentile value.

### Statistical Analysis

Unless otherwise specified, quantitative data were compared as follows: for pairwise comparisons between two independent groups, two-tailed Mann-Whitney U tests (Wilcoxon rank-sum tests) were performed. For group-wise comparisons involving three or more independent groups, one-way Analysis of Variance (ANOVA) was conducted to control for family-wise type I error rate arising from multiple testing, raw *p*-values across all primary outcome tests were adjusted using the Benjamini-Hochberg False Discovery Rate (FDR) procedure. Adjusted *p*-values *<* 0.05 were considered statistically significant. All statistical analyses were implemented in Python (v3.12) using scipy.stats and statsmodels libraries. All raincloud plots (e.g. 1D, G, 3I, J, etc.) show median (open circle), Q1-Q3 (thick vertical bar), kernel density estimation with half violin (left) and individual observations as scatter points (right).

### Single-cell RNA sequencing

Embryos were harvested and electroporated with pCAG-GFP to tag the node endoderm (NE) at stages HH8, HH11, and HH11, then developed in EC culture to HH10, HH15, and HH18, respectively, as described above. Tailbud tissues were dissected in cold PBS, removing the extraembryonic membranes. 12-15 embryos were pooled per time group. Cells were enzymatically dissociated in Accutase (STEMCELL Technologies), cell viability (>90%) and concentration was assessed with a Countess 3 automated cell counter with AO/PI staining. Single-cell libraries were generated using the Chromium GEM-X Single Cell 3’ Reagent Kits v4 (10x Genomics, Pleasanton, CA) according to the manufacturer’s instructions, aiming for approximately 20,000 targeted cell recovery per batch (time point). Resulting libraries were amplified using dual-index primers (Dual Index Kit TT Set A), then pooled and sequenced on an Illumina NovaSeq X Plus platform to a sequencing depth of approximately 50,000 mean reads per cell.

Raw sequencing data (FASTQ files) were processed using the Cell Ranger 9.0 pipeline (10x Genomics), aligning reads to the Gallus gallus reference transcriptome (GRCg7b), to produce the counts matrix. Down-stream analysis was performed using Scanpy (v1.12) (*73*). Low-quality cells were excluded based on the following criteria: (1) cells with too few (e.g. *<*200) or too many (e.g. >8,000) detected genes; (2) cells with more than 10% of total UMI counts originating from mitochondrial genes; and (3) cells with fewer than 1,000 total UMIs, following sc-best-practices recommendations (*74*). Doublets were predicted and removed using Scrublet (*75*). Gene expression matrices were normalized to 10,000 counts per cell and log-transformed. Highly variable genes (HVGs) were identified using the highly_variable_genes function (flavor=’seurat’) (*74*). Principal Component Analysis was performed on the scaled data, and the first 50 principal components were used to construct a neighborhood graph. Dimensionality reduction was performed by Pairwise Controlled Manifold Approximation (PaCMAP) (*76*). To recover GFP transcripts in the dataset, a corresponding gene sequence was added in the gene annotation file (GTF), and GFP+ counts threshold was determined by inspecting label persistence per batch (Supplemental Fig.S2B). To define cell populations, we performed unsupervised Leiden clustering, with sweeping resolutions until over-clustering, then merging into cell types by scoring on predefined cell marker scores (Supplementary Table S1); for this list, we consulted published data from NMP single-cell studies (*6*) and chick datasets(*3*, *7*). To investigate the developmental trajectories revealed by the data we employed the CellRank2 package (*55*) to reconstruct cell state dynamics and infer cellular trajectories. We employed the the RealTimeKernel (optimal transport based matching of experimental time points) and a precomputed PseudotimeKernel, using the CytoTRACE2 potency score(*53*), and combined the two kernels in equal weight for the computation of fate probabilities. Subsequently, we defined terminal lineages using the reference macrostates identified from CellRank. To assemble unique trajectories from NE vs NMP populations, we subset the data accordingly to compute appropriate lineage absorption scores and respective lineage drivers (genes that maximally correlate with given lineage or trajectory).

### TrackerSeq

The TrackerSeq plasmid library was generated as described in detail previously (*54*). Electroporation of the TrackerSeq plasmid solution (1.0 µg/µL helper and 1.0 µg/µL donor library) was performed *ex ovo* at HH10 and embryos grown to HH18 in EC culture as described above. After tissue dissection and single-cell dissociation, GFP+ cells were enriched by FACS (Columbia Stem Cell Initiative Flow Core) on a Bigfoot 50204 (Thermo Fisher) cytometer, and the NovoExpress 1.6.2 software. Sorted cells were processed for single-cell sequencing with the GEM-X (10X Genomics) protocol as above, with separate libraries created for transcriptome and barcodes (*54*). TrackerSeq barcode reads were processed following (*54*). Briefly, R2 FASTQ files were trimmed to remove sequence flanking the lineage barcodes (BC), and cell barcodes and UMIs extracted from R1 reads were appended to the corresponding BC read names. The resulting reads were converted into a sparse matrix with cell barcodes as rows and BCs as columns. CloneIDs were assigned by hierarchically clustering cell barcodes based on Jaccard similarity using scikit-learn (Python), and the resulting dendrogram was cut by visual inspection to obtain stable clonal groupings.

## Footnotes

## Acknowledgments

We thank members of the Nerurkar lab for their valuable scientific input, and in particular Nathalie Houssin, and Olivia Powell for help with mRNA electroporations. We also thank the McFaline-Figueroa lab and Tosches lab members for help with single-cell experiments and TrackerSeq, the CSCI Flow Core (Michael Kissner) for the FACS experiments and Charlene Guillot for critically discussing the results.

## Author Contributions

Conceptualization: P.O., L.C., C.M., J.M.F., N.L.N.; Methodology: P.O., L.C., G.G., C.L., L.S., N.L.N.; Formal analysis: P.O., L.C., D.D., J.Y.P., N.L.N.; Investigation: P.O., L.C., D.D., J.Y.P., N.L.N.; Resources: N.L.N.; Data curation: P.O., L.C., D.D., J.Y.P., N.L.N.; Writing – original draft: P.O., N.L.N.; Writing – review & editing: P.O., L.C., N.L.N.; Supervision: N.L.N.; Funding acquisition: N.L.N.

## Funding

This work was funded by the NIGMS (R35GM142995, N.L.N.) with additional support from the Columbia University Digestive and Liver Disease Research Center (1P30DK132710).

## Conflict of Interest

The authors have no conflict of interest to declare.

## Data and Code Availability

scRNA-seq data will be deposited on GEO. Code associated with the analyses performed can be found at [Github](https://github.com/eigenP/OikonomouNerurkar2026).

## SUPPLEMENTARY INFORMATION

**Figure S1.**
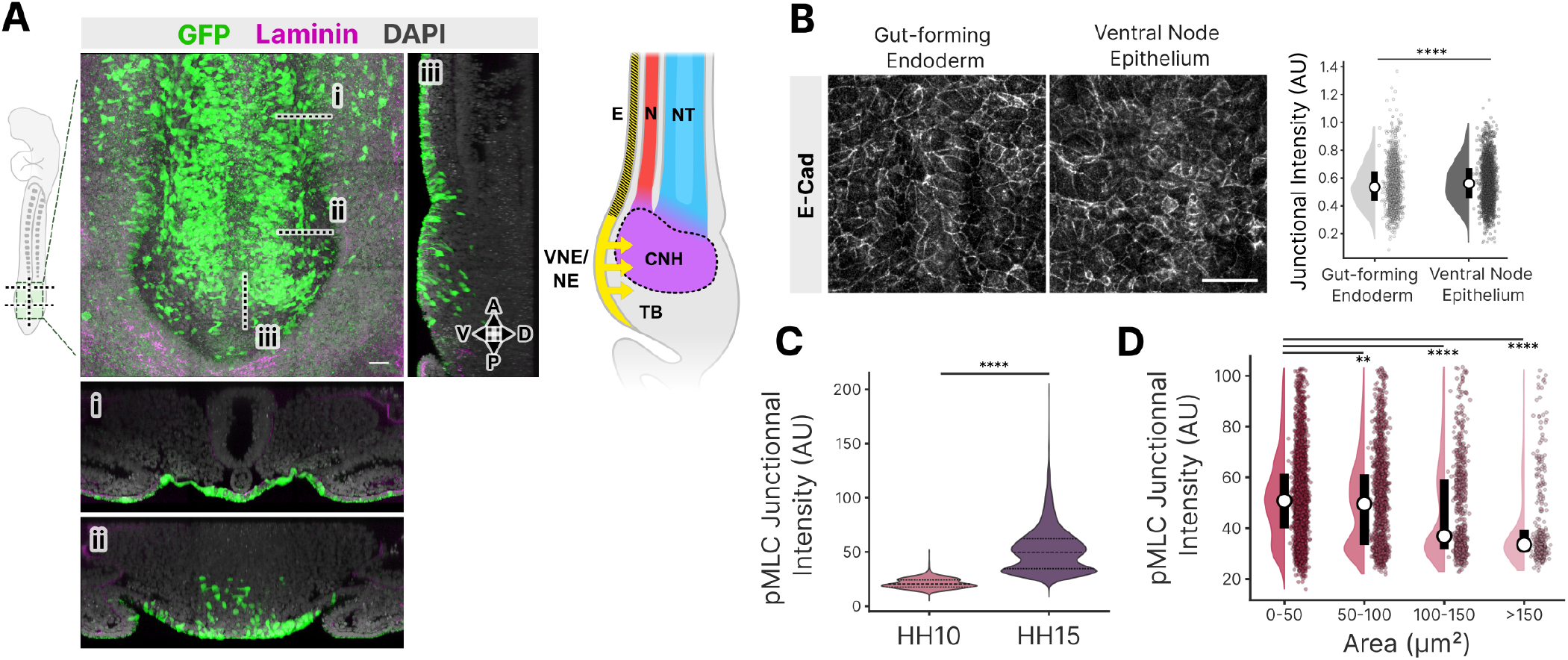
Ingression of ventral node epithelium coincides with apical myosin activity. (A) Ventral view of an optically cleared HH15 embryo following electroporation with pCAG-GFP at HH10 to label the ventral epithelium of the posterior embryo, with i and ii indicating transverse views anterior to the node and within the node region, respectively, visualized below, and iii indicating the saggital view at right. Sample immunostained for GFP (green) and laminin (magenta), counterstained with DAPI (gray); Scale = 20µm. Schematic at right indicates sagittal view of node cell ingression from the ventral node epithelium (VNE), later identified as node endoderm (NE) into the chordoneural hinge (CNH) region of the tailbud (TB), which lacks the planar organization of germ layers that is apparent anterior to the node, with distinct demarcation of definitive endoderm (E), notochord (N), and neural tube (NT). Anterior (A), Posterior (P), Ventral (V), and Dorsal (D) axes are as indicated in (iii). (B) Immunostaining for E-Cadherin in gut-forming endoderm and the ventral node epithelium at HH10, quantified at right (N = 6 embryos; n = 258 (node endoderm) and 180 (gut forming endoderm) cells; Scale = 20µm. (C) Quantification of junctional intensity of pMLC from immunostaining at stage HH10 and 15. (D) Data from C binned according to cell apical area at HH15. ** p *<* 0.01, **** p *<* 0.0001.

**Figure S2.**
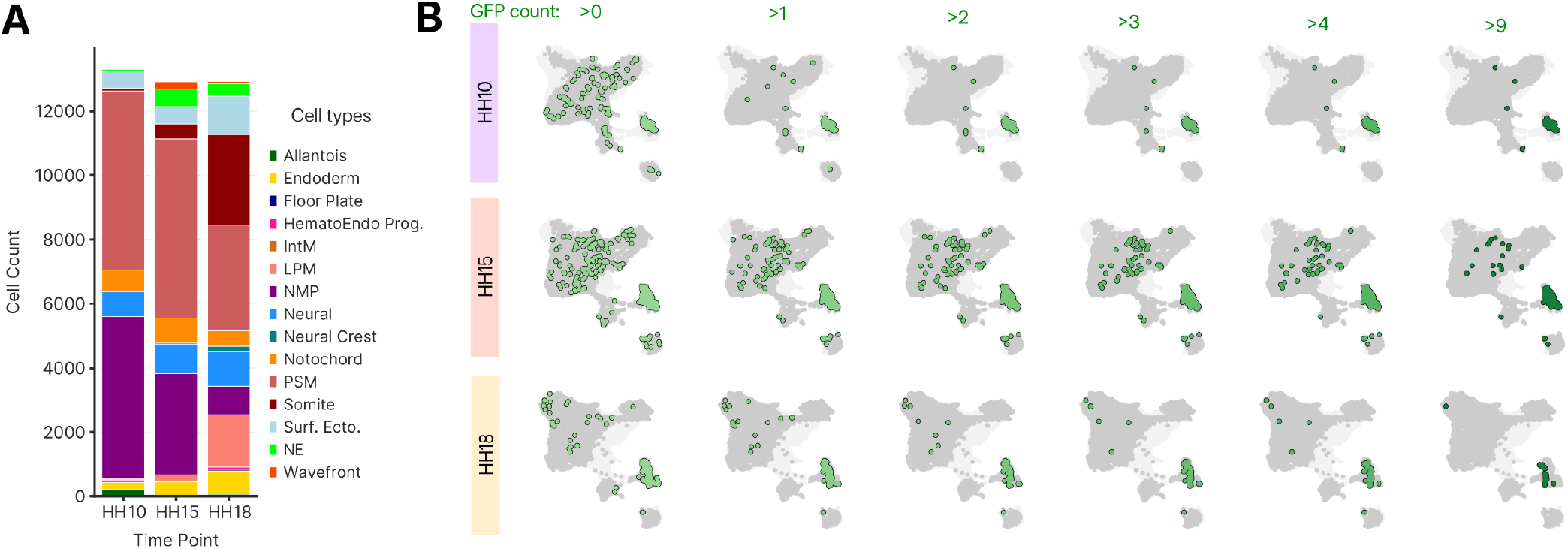
scRNA-seq of the posterior chick embryo following electroporation-based tagging of VNE and its descendants. (A) Cell-type composition of the single-cell RNA-sequencing dataset at HH10, HH15, and HH18. (B) Thresholding of GFP transcript counts to determine GFP+ cells at HH10, HH15, and HH18, with cells above threshold indicated in green and overlaid on PaCMAP embedding of stage-integrated dataset, where GFP-negative cells of the corresponding developmental stage are indicated in dark gray, and cells from other time points in light gray.

**Figure S3.**
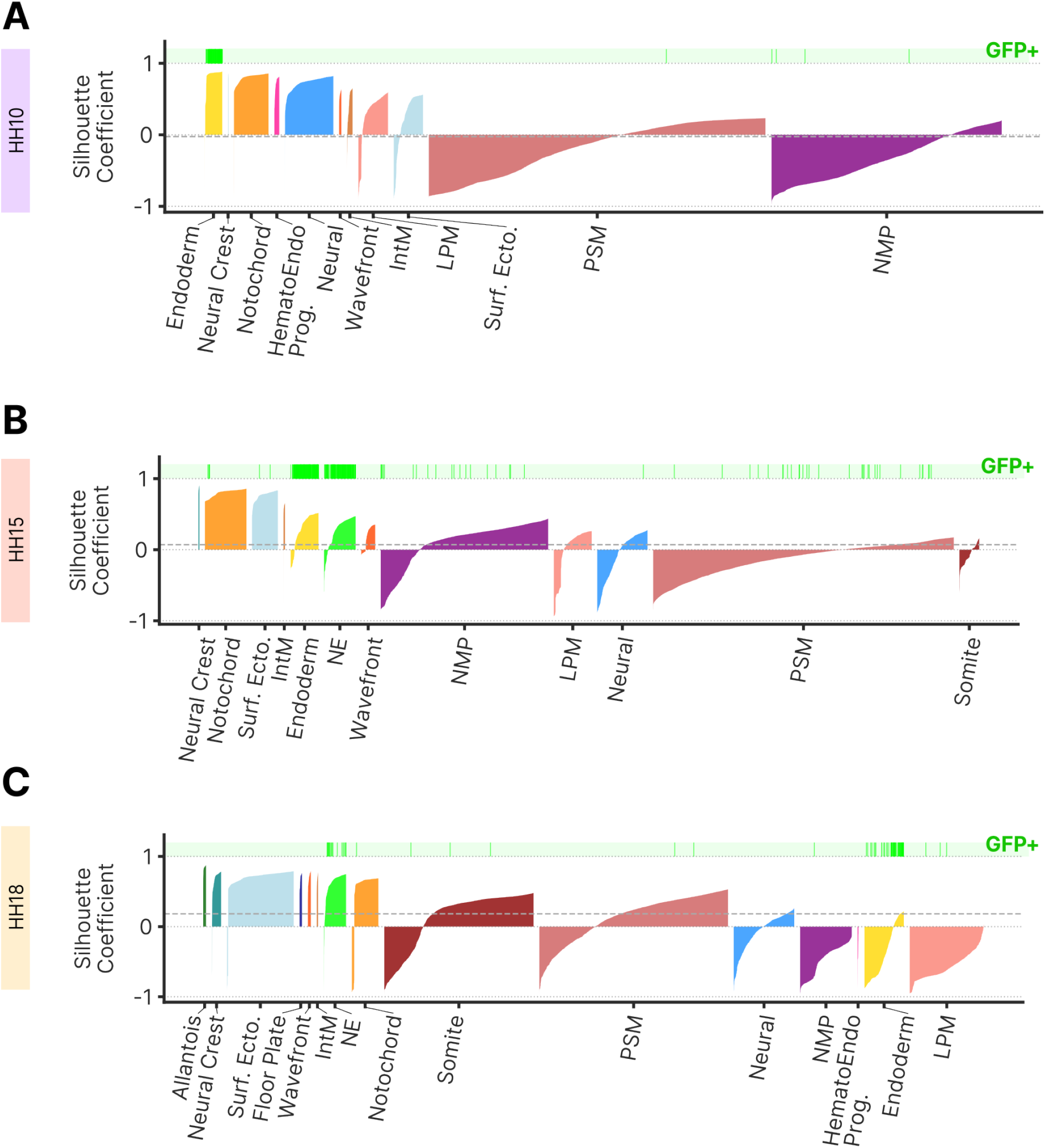
Cell type cohesion and separation. (A–C) Silhouette coefficients for cells by cell type at (A)HH10, (B)HH15, and (C)HH18. Green hash marks over each plot indicate position of each GFP+ cell detected.

**Figure S4.**
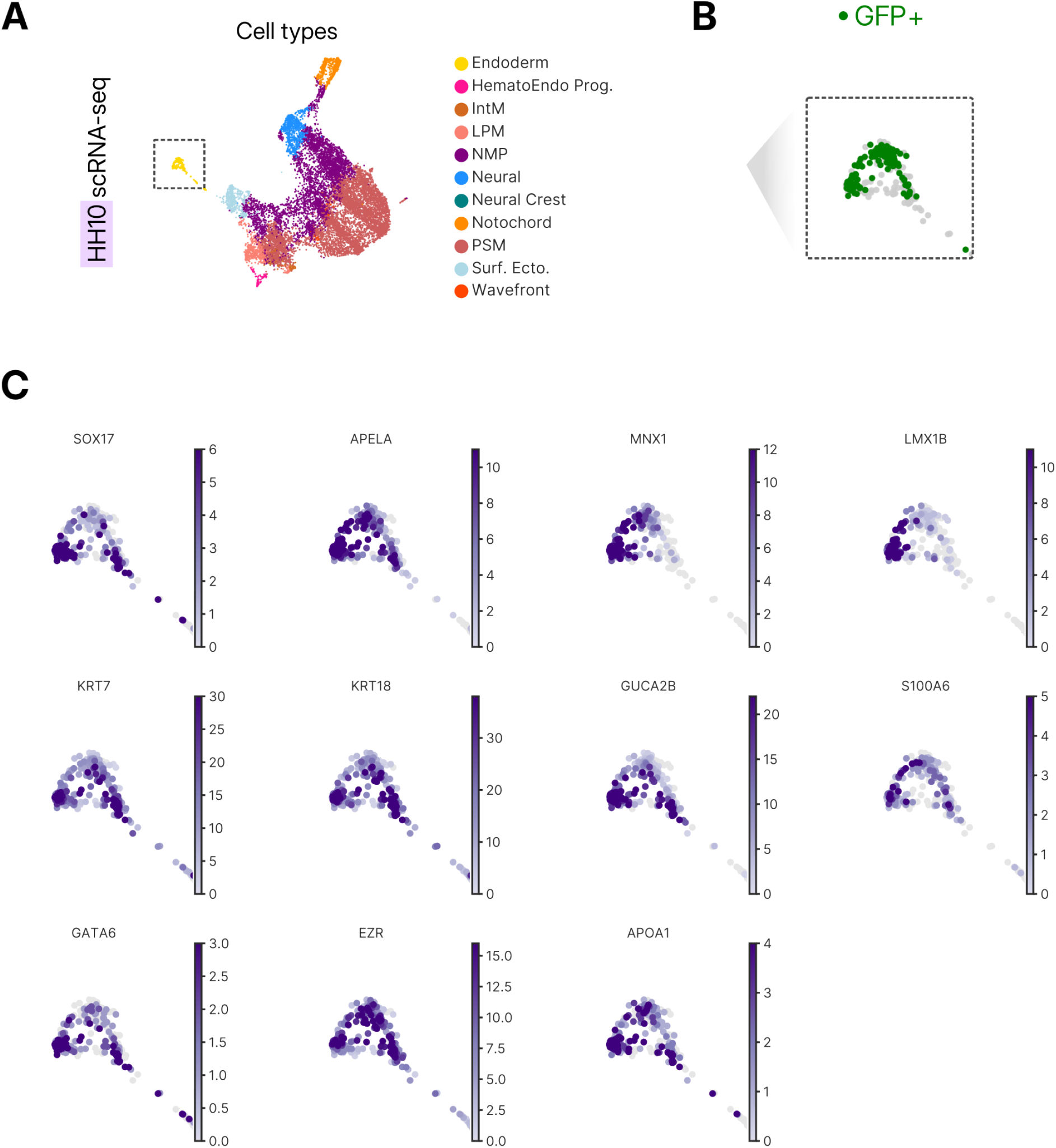
Expression of endoderm-associated genes in the HH10 endoderm cluster. (A) PaCMAP embedding of the HH10 scRNA-seq dataset (data from Fig. 2F). The dashed box indicates the endoderm cluster magnified in (B, C). (B) Distribution of GFP+ cells within the HH10 endoderm cluster. (C) Expression of top differentially expressed genes enriched in the endoderm cluster when compared to all other cell types at HH10.

**Figure S5.**
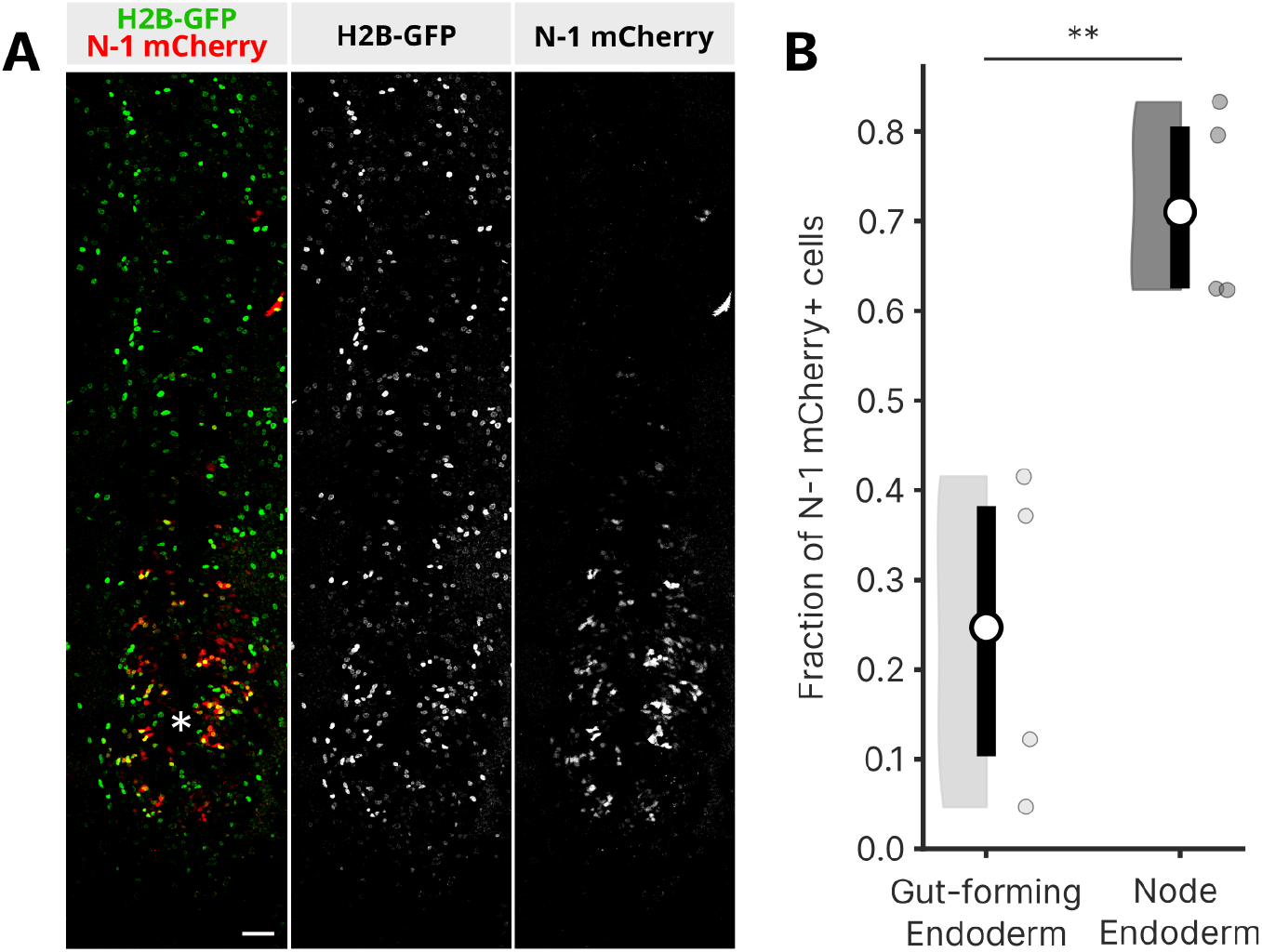
N-1 enhancer of SOX2 is active in posterior endoderm. (A) Ventral view of HH15 embryo co-electroporated with N-1 mCherry reporter and pCAG-H2B-GFP at HH10; * denotes location of Hensen’s node; Scale = 50µm. (B) Fraction of electroporated cells (GFP+) that have N-1 enhancer activity (mCherry+) in gut-forming and node endoderm; Paired t-test, ** p < 0.01.

**Figure S6.**
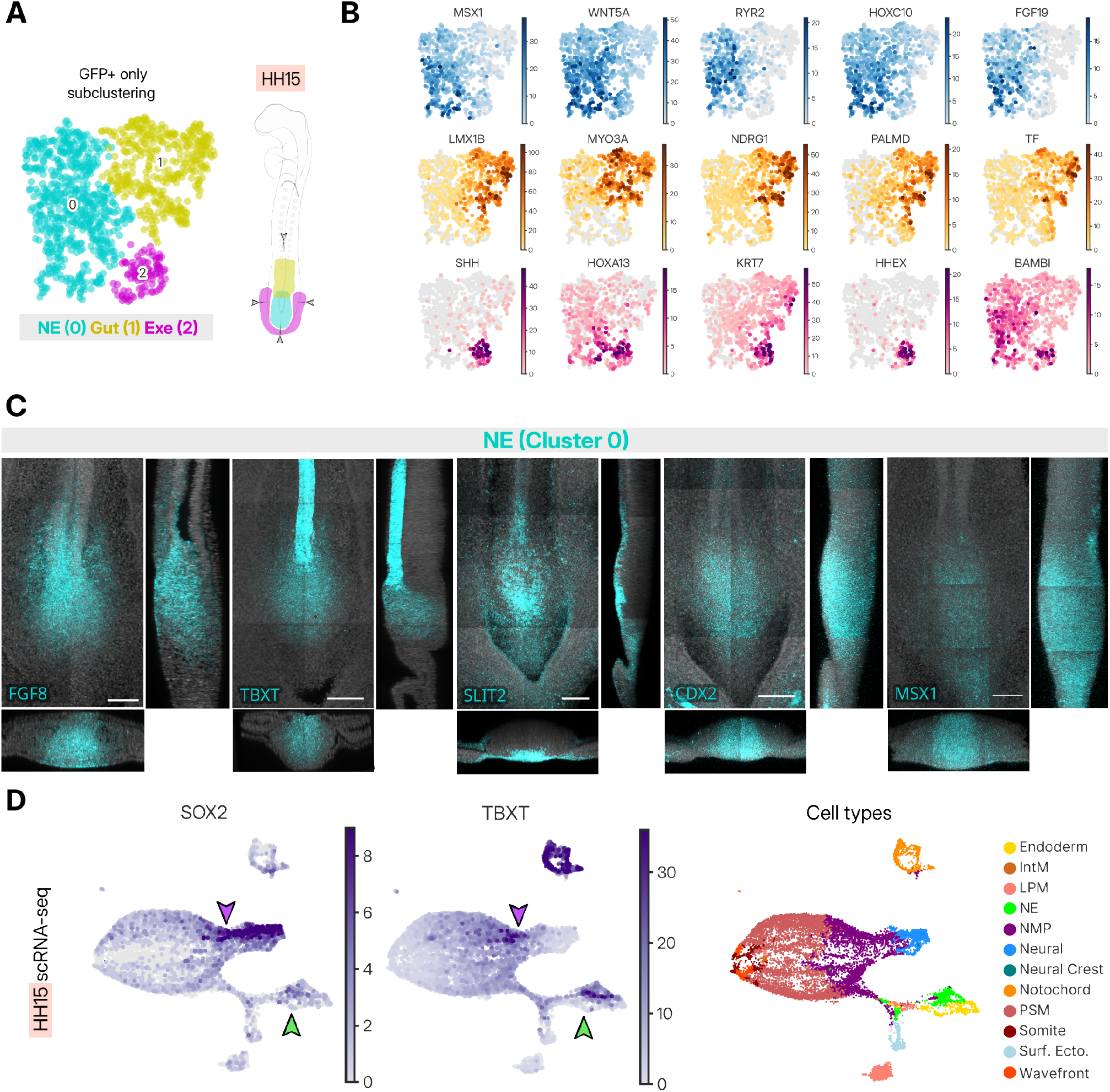
Identification of distinct endodermal subpopulations in the posterior HH15 embryo. (A) PaCMAP subclustering of HH15 endoderm with three distinct cluster identities identified: NE (node endoderm, cyan), gut forming endoderm (Gut, yellow), and extraembryonic endoderm (ExE, magenta). Schematic represents proposed domains of each subtype, with arrowheads indicating orthogonal views in (C). (B) Top differentially expressed genes associated with each of the three subclusters in (A). (C) HCR validation of candidate marker genes for the NE population; Scale = 100µm. (D) PaCMAP embedding of SOX2 and TBXT expression at HH15, with co-expression in NMP and NE clusters indicated by purple and green arrows, respectively.

**Figure S7.**
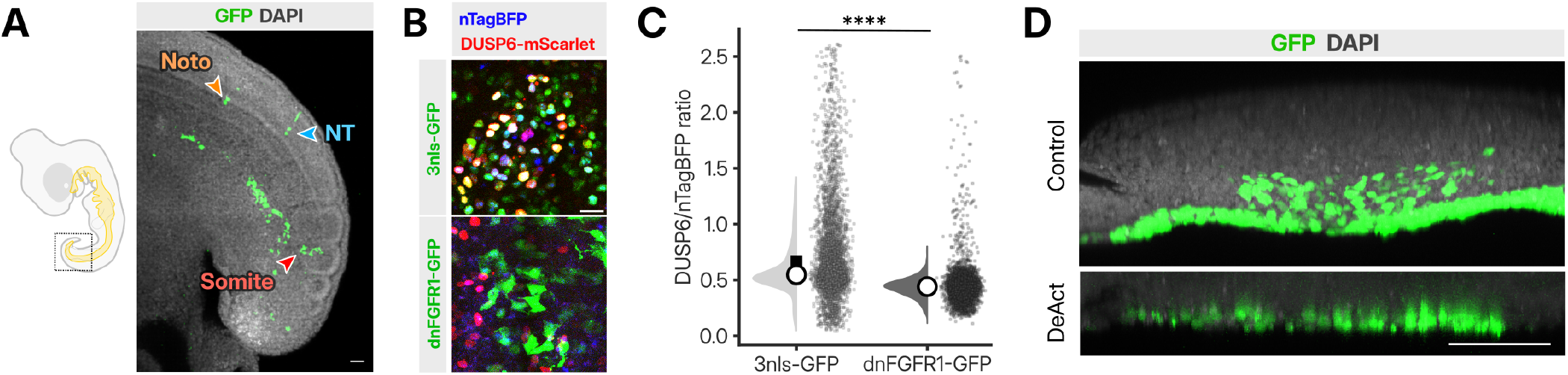
Fate mapping and FGF inhibition of the NE. (A) GFP+ cells appearing in the notochord (orange arrow), neural tube (blue arrow), and somite (red arrow). (B) Representative images of embryos electroporated with 3nls-GFP and dnFGFR1-GFP plasmid. (C) Quantification of DUSP6 to nTAGBFP ratio, comparing embryos electroporated with 3nls-GFP (N = 4 embryo; n = 6214 cells) and dnFGFR1-GFP (N = 4 embryos; n = 5200 cells) in (B). **** p *<* 0.0001. (D) Transverse view of HH15 embryos, comparing ingression between control (pCAG-GFP) and DeAct electroporated samples.

**Figure S8.**
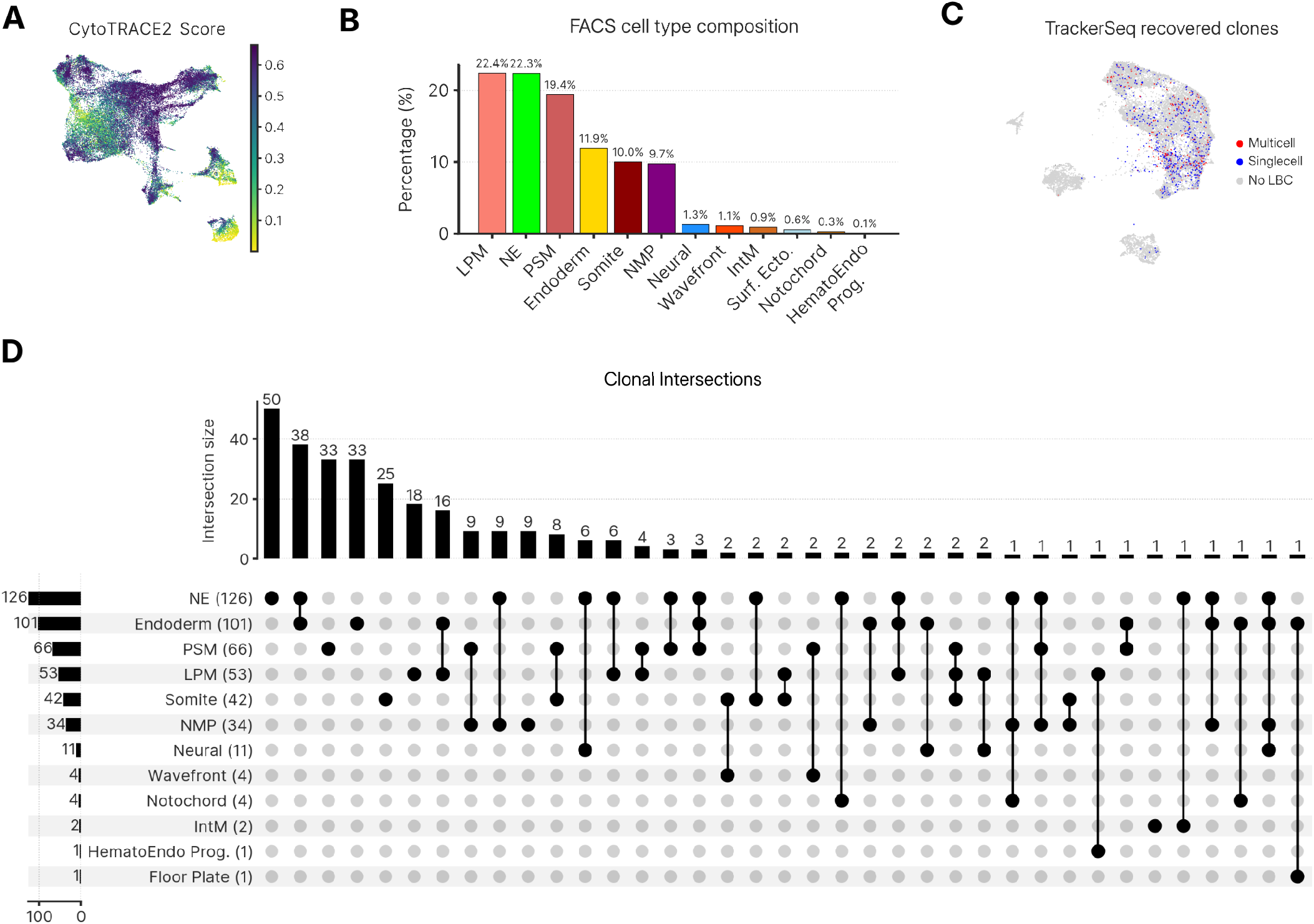
Multipotency of the node endoderm. (A) CytoTRACE2 potency score, absolute continuous scale (1 totipotent, 0 terminally differentiated). (B) Distribution of TrackerSeq cells with recovered lineage barcodes, classified as multicell clones, single-cell clones, or cells without a recovered lineage barcode (LBC). (C) Cell-type composition of the FACS-isolated population. (D) UpSet plot showing the number and cell-type composition of recovered multicell clones. Connected dots indicate cell types represented within the same clone. Bars indicate the number of clones corresponding to each intersection.

**Figure S9.**
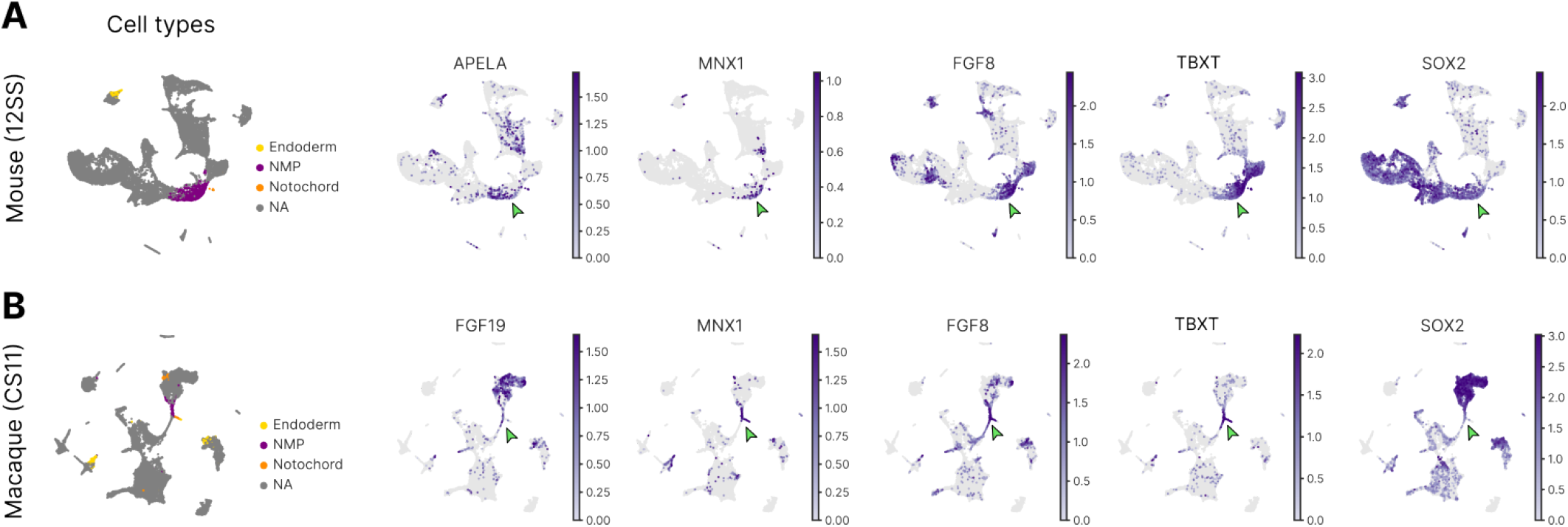
Gene signature of node endoderm identified in NMP population of published mouse and macaque scRNA-seq datasets. (A) PaCMAP embeddings of a 12-somite-stage mouse scRNA-seq dataset showing annotated endoderm, NMP, and notochord populations, followed by the expression of *APELA*, *MNX1*, *FGF8*, *TBXT*, and *SOX2*. (B) PaCMAP embeddings of a Carnegie stage 11 macaque scRNA-seq dataset showing the corresponding cell populations, followed by the expression of *FGF19*, *MNX1*, *FGF8*, *TBXT*, and *SOX2*. Green arrowheads indicate the putative NE population.

**Table S1.** Genes used for cell type annotation in the scRNA-seq dataset.

| **Cell Type** | **Marker Genes** |
| --- | --- |
| Endoderm | *APELA, MNX1, S100A6, BAMBI, GUCA2B, KRT18, EPCAM* |
| Neural | *HES5, PAX6, RFX4, GBX2, SOX2, SOX1, ZIC2, NES* |
| Surf. Ecto. | *PDGFA, DSP, CDH1, WNT6, IRF6, TFAP2A, KRT8, KRT19* |
| Neural Crest | *SOX10, SOX9, PAX7, FOXD3, ZIC1* |
| Floor Plate | *NKX6-2, FOXA2, SHH, NKX2-2, NKX2-9, APELA* |
| PSM | *MSGN1, TBX6, HES7, LFNG* |
| Wavefront | *MESP2, RIPPLY1, RIPPLY2, EPHA4, CDH20, DLL1* |
| Somite | *TWIST1, TCF15, FOXC2, PAX1* |
| LPM | *PRRX1, BMP4, NR2F2* |
| IntM | *PAX8, OSR1, WT1, MEIS2* |
| Notochord | *NOTO, SHH, TF, CHRD, LEFTY1, SLIT3, TBXT* |
| NMP | *WNT3A, NKX1-2, WNT8A, SOX2, TBXT, CDX2, CDX4, CYP26A1, EPHA1, FGF8, FGF19* |
| NE | *APELA, MSX1, FGF19, MNX1, FGF8, EPCAM* |

## Notes

### Competing Interest Statement

The authors have declared no competing interest.

### Summary of Updates

This revised preprint adds considerable scientific content, to extend and expand on the premise of the original preprint, including a deeper characterization of the epithelial to mesenchymal transition of node endoderm, additional data analysis and new experiments supporting the endodermal identity of cells lining the ventral surface of Hensen's node, new results supporting the transition of cells from endodermal to neuromesodermal fates that persist later in development, and a deeper investigation of inferred fate convergence dynamics. In addition, the revision includes 9 new supplemental figures that clarify key points of the work, including methodology and new findings.

